# eNetXplorer: an R package for the quantitative exploration of elastic net families for generalized linear models

**DOI:** 10.1101/305870

**Authors:** Julián Candia, John S. Tsang

## Abstract

**Background:** Regularized generalized linear models (GLMs) are popular regression methods in bioinformatics, particularly useful in scenarios with fewer observations than parameters/features or when many of the features are correlated. In both ridge and lasso regularization, feature shrinkage is controlled by a penalty parameter λ. The elastic net introduces a mixing parameter *α* to tune the shrinkage continuously from ridge to lasso. Selecting *α* objectively and determining which features contributed significantly to prediction after model fitting remain a practical challenge given the paucity of available software to evaluate performance and statistical significance.

**Results:** eNetXplorer builds on top of glmnet to address the above issues for linear (Gaussian), binomial (logistic), and multinomial GLMs. It provides new functionalities to empower practical applications by using a cross validation framework that assesses the predictive performance and statistical significance of a family of elastic net models (as *α* is varied) and of the corresponding features that contribute to prediction. The user can select which quality metrics to use to quantify the concordance between predicted and observed values, with defaults provided for each GLM. Statistical significance for each model (as defined by *α*) is determined based on comparison to a set of null models generated by random permutations of the response; the same permutation-based approach is used to evaluate the significance of individual features. In the analysis of large and complex biological datasets, such as transcriptomic and proteomic data, eNetXplorer provides summary statistics, output tables, and visualizations to help assess which subset(s) of features have predictive value for a set of response measurements, and to what extent those subset(s) of features can be expanded or reduced via regularization.

**Conclusions:** This package presents a framework and software for exploratory data analysis and visualization. By making regularized GLMs more accessible and interpretable, eNetXplorer guides the process to generate hypotheses based on features significantly associated with biological phenotypes of interest, e.g. to identify biomarkers for therapeutic responsiveness. eNetXplorer is also generally applicable to any research area that may benefit from predictive modeling and feature identification using regularized GLMs.

**Availability and implementation:** The package is available under GPL-3 license at the CRAN repository, https://CRAN.R-project.org/package=eNetXplorer

## Background

Rigorous, exploratory analysis for the identification of correlates and predictors in a multi-parameter/feature setting is needed in a variety of contexts, especially in systems biology where data involving a large number of parameters are highly prevalent. Oftentimes, bioinformatics analysis in such settings involves generalized linear models where observations (*N*) are outnumbered by parameters/features (*p*) measured. This class of problems can be addressed by the elastic net, ^1^ which uses a mixing parameter *α* to tune the number of features used in the model continuously from ridge (*α* = 0) to lasso (*α* = 1).

Algorithmically, the elastic net was efficiently implemented by the package glmnet, a coordinate descent algorithm ^2,3^ that, for each *α*, generates an entire path of solutions in the regularization parameter λ, which controls the penalty for using more parameters. While the choice of λ is usually guided by prediction performance using cross validation, *α* is often viewed as a higher-level parameter and chosen based on more subjective grounds. ^3^ Lasso generates parsimonious solutions in that a small number of predictor variables are selected from a large number of input parameters, particularly useful in *p* ≫ *N* scenarios; however, in the presence of complex correlation structures among input variables (or degeneracies), lasso can arbitrarily pick one as a predictor among a set of correlated variables and ignore the rest. This characteristic may lead to models that are idiosyncratic of the input data set, as opposed to more robust solutions capturing relevant signals, or it may even lead to unstable solutions in some extreme cases. ^3^ On the contrary, ridge regression promotes redundancy by shrinking correlated features towards each other, thus allowing information to be borrowed across them.

In multi-parameter exploratory analysis where the primary goal is to generate hypotheses, e.g. to assess which variables correlate with a biological phenotype of interest, it is desirable to examine the entire family of elastic net models spanning the range from ridge to lasso. In this scenario, an objective, quantitative framework is needed to assess the statistical significance of individual models and, within each model, that of individual parameters/features. Towards this goal of transforming large-scale data sets into biological hypotheses, this paper introduces eNetXplorer, an R package providing a quantitative framework to explore elastic net families for generalized linear models (GLM). In the current version, three important GLM types are implemented: linear regression, two-class logistic, and multinomial classification. In future releases, we plan to extend it to other GLM types such as Poisson regression and the Cox model for survival data.

eNetXplorer is built on top of the existing R package glmnet and provides new functionalities to empower practical applications, including evaluating the statistical significance of a family of fitted models and the corresponding features that contributed significantly to prediction via a cross validation framework. Both bioinformaticians and biologists can utilize our package to help transform data into biological insights, for example, to help answer which biological variables, often out of a large number in the current age of large-scale ‘omics’ datasets, provide predictive information about an outcome variable (e.g., drug responses). Furthermore, our package provides a set of standard plots, summary statistics, and output tables to enable the visualization and interpretation of the results, thus making regularized GLMs more readily accessible to a larger user base of diverse scientific backgrounds.

Fig. 1 provides a conceptual schema of eNetXplorer in the context of GLM regularization. Multiple datasets of size *N* × *p*_*d*_ (*d* = 1, …, *D*) can be aggregated into an input matrix *N* × *p*, where 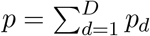 (Fig. 1(a)). In *p* ≫ *N* scenarios, such as (but not limited to) typical single- and multi-omics datasets, regression analysis requires regularization models such as ridge and lasso (Fig. 1(b)). The elastic net provides an integrated framework to analyze the full regularization path from ridge to lasso; however, there remained a number of issues ranging from model selection and assessing statistical significance of individual models to feature selection and their statistical significance (Fig. 1(c)). By generating null-model ensembles via random permutations of the sample label of the response variable (Fig. 1(*d*_1_)), eNetXplorer addresses these issues (Fig. 1(*d*_2–4_)). Although the emphasis of our presentation is placed on biomedical applications, for which high-throughput technologies such as DNA/RNA sequencing, deep-phenotyping flow and mass cytometry, as well as highly multiplexed proteomics typically generate *p* ≫ *N* datasets, eNetXplorer is generally applicable to datasets beyond biomedicine. The accompanying vignette (Additional file 1) illustrates in detail the application of eNetXplorer to synthetic datasets with different feature/response co-variance structures, which further highlight the flexibility of our approach in a variety of scenarios.

**Figure 1:**
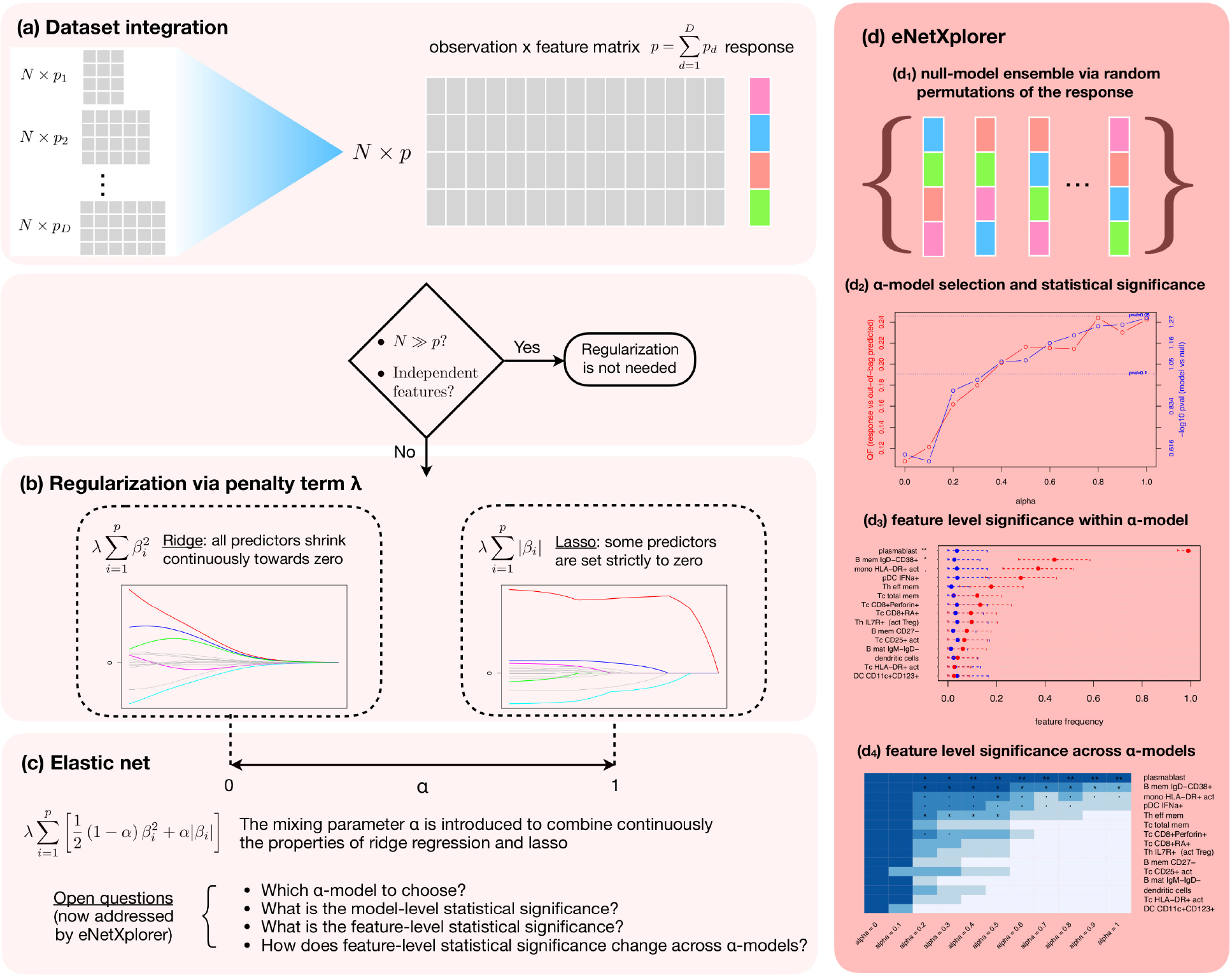
Conceptual schema of eNetXplorer. (a) One or more datasets can be aggregated into an *N* × *p* input matrix. Depending on the GLM of interest, the response is a numeric vector (linear), 2-class factor (binomial) or multi-class factor (multinomial). (b) Ridge and lasso implement different regularization penalty terms, which are tuned by the regularization parameter λ. (c) The elastic net introduces the mixing parameter *α* as a continuous tuner from ridge to lasso. (d) eNetXplorer generates a null-model ensemble via random permutations of the response (*d*_1_), which allows the quantitative exploration of elastic net families by assessing the statistical significance of each model (*d*_2_), the feature-level significance within each model (*d*_3_) and across the entire elastic net family (*d*_4_).

## Implementation

eNetXplorer generates a family of elastic net models for multiple values of *α* from ridge (*α* = 0) to lasso (*α* = 1). Fig. 2 shows a flowchart of the algorithm’s implementation. The algorithm is composed of three main modules: (a) model building, (b) null model building, and (c) model vs null comparison, which are sequentially executed for each value of *α*; at the end, the results are integrated across *α* for downstream analysis and visualization.

**Figure 2:**
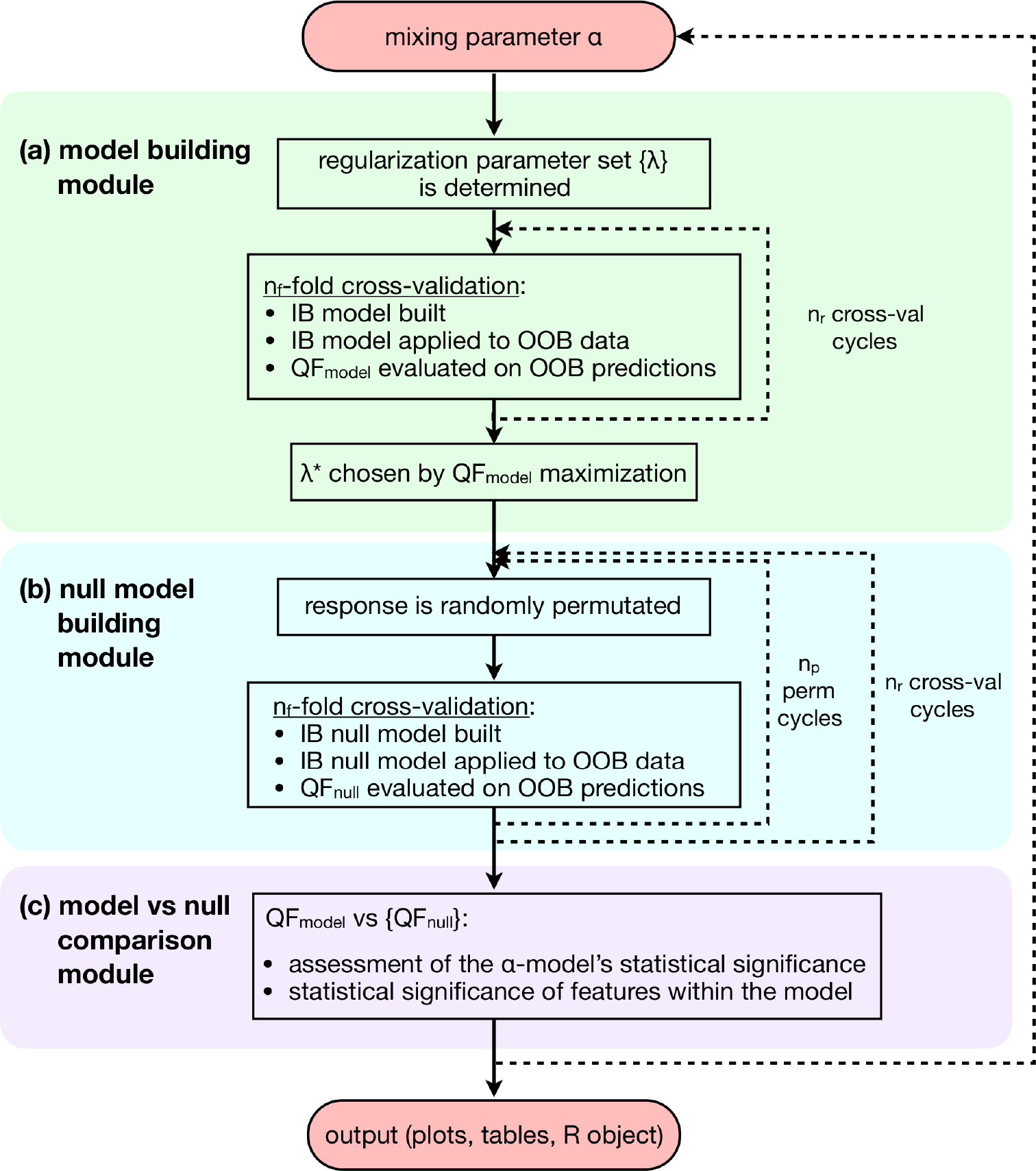
Flowchart of the algorithm’s implementation. The algorithm consists of three main modules: (a) model building, (b) null model building, and (c) model vs null comparison, sequentially executed for each value of *α*; at the end, the results are integrated across *α* for downstream analysis and visualization. Abbreviations used: in-bag (IB), out-of-bag (OOB), quality function (QF). More details provided in the Implementation Section.

In the model building module (Fig. 2(a)), a set of *n*_λ_ values is obtained using the full data; independently from *n*_λ_, the user may also specify a value for 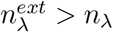 to extend the range of λ values symmetrically while keeping its density constant in log scale. For each λ, elastic net cross-validation models are generated for *n*_*r*_ runs, where each run randomly assigns instances (i.e. the *N* measured samples/observations) among *n*_*f*_ folds. The chosen regularization λ* is determined by maximizing a quality function (QF) that compares the out-of-bag (OOB, i.e. not used in training) predicted response against the observed response. User-defined QFs can be provided. Otherwise, GLM-specific defaults are used: for linear regression, the default QF is correlation (where the user can choose among Pearson’s, Spearman’s and Kendall’s methods); for binomial models, it is accuracy; and, for multinomial models, average accuracy. For the latter two, the QF defaults are chosen based on the property of invariance under class label permutations. Other popular performance measures implemented are precision, recall (sensitivity), F-score, specificity, and area-under-the-curve, which are not invariant under class label permutations, ^4^ but may be useful for some applications. Any of these performance measures can be selected as QF by the end user.

Individual features are characterized by their distribution of model coefficients across cross-validation iterations, which we summarize by the following measures. From the *feature frequency* per run, 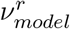, defined as the fraction of folds (within a run) for which the given feature was assigned a non-zero model coefficient, we derive the mean and standard deviation of feature frequency (averaged over all runs). Similarly, we define the *feature coefficient* per run, 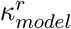 as the mean of non-zero model coefficients across all folds in the run. We perform a weighted average over all runs, where weights are 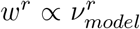, to determine the weighted mean and weighted standard deviation of the feature coefficient.

A key aspect of eNetXplorer is the generation of an ensemble of null models associated with each (*α*–specific) member of the elastic net model family, which is accomplished by the null model building module (Fig. 2(b)). Each one of *n*_*r*_ runs are assigned into folds (based on the same fold assignments used previously) and *n*_*p*_ null models per run are generated by randomly shuffling the sample labels of the response; for each permutation, the overall OOB performance of the null model is evaluated via the QF, whereas the contribution of individual features is characterized by 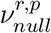 and 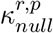, following analogous definitions to those given above.

The empirical statistical significance of a model, implemented by the model vs null comparison module (Fig. 2(c)), is hence determined as

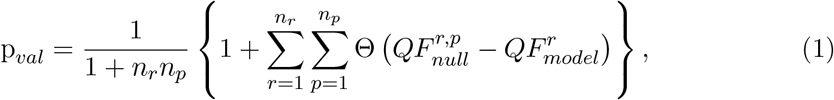

where Θ is the right-continuous Heaviside step function. For sampling permutations with replacement, this expression provides a conservative estimate; ^5^ expressions for the exact p-value, as well as numerical approximations thereof, are provided by Ref. ^6^

As discussed above (recall Fig. 1), eNetXplorer aims to tackle the following questions that remained unaddressed by the elastic net framework implemented in glmnet:

- Which *α* provides the best-performing regularized model?
- What is the statistical significance of the predictive performance for each model across different *α*?
- For each *α*, what is the statistical significance, in terms of contribution to prediction, of individual features included in the model? And how does the statistical significance change across *α*?

In order to address these questions, eNetXplorer generates quantitative results and provides a variety of standard plots (Figs. 3-6) that enable their interpretation.

**Figure 3:**
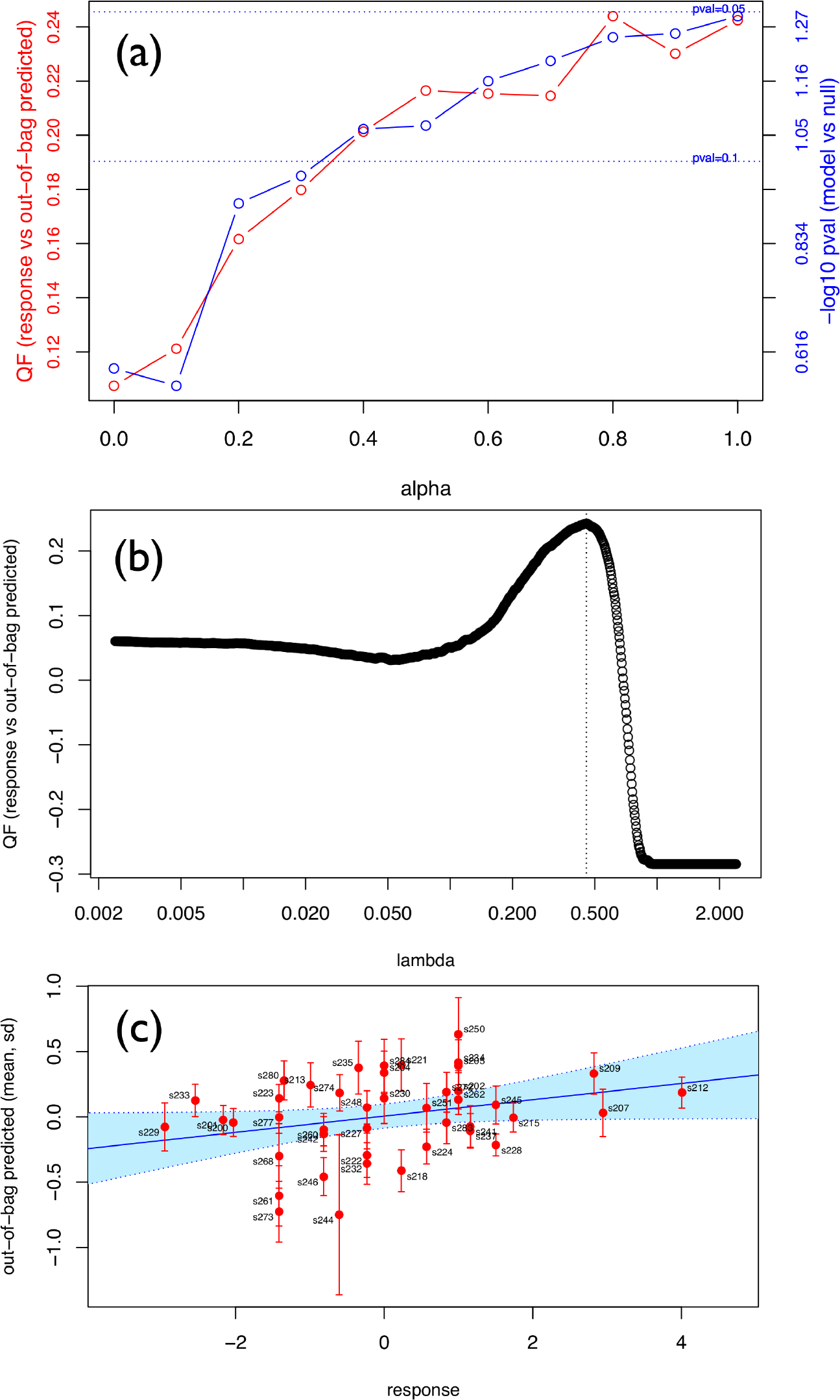
Linear regression model of postvaccination immune cell frequency correlates of H1N1 titer response. (a) Performance across *α*: quality function (QF) and statistical significance (p-value of model vs null). The default QF for linear regression is Pearson’s correlation between out-of-bag predictions and the response. (b) Selection of λ by QF maximization (*α* = 1). (c) Scatterplot of response vs out-of-bag predictions across all subjects (*α* = 1); best linear fit (solid line) and 95% CL (shaded region) also shown.

Let us highlight a few key graphical outputs of our package, generated by the use case studies below, wherein further details on the data and goals of the analyses can be found. Model performance results are visualized by a summary plot, which shows the average OOB QF (red plot, left axis) and the model vs null p-value significance (blue plot, right axis) spanning the full range of *α* values for the example discussed below (Fig. 3(a)); Fig. 3(b) shows the lasso QF vs λ profile and the chosen λ* (dashed line); using this λ*, Fig. 3(c) shows OOB predictions vs response for all observations in the dataset.

Replacing QF in Eq. (1) by *ν* or |*κ*|, our framework also provides empirical p-value estimates of the importance and statistical significance of individual features. Caterpillar plots are generated to display the top features ranked by their importance, in which significance thresholds are indicated by customary dot and asterisk annotations. Fig. 4(a) shows the top features ranked according to statistical significance based on the frequency at which each feature is selected across cross validation iterations; red symbols and bars represent the mean and standard deviation of the model feature frequency, while those for the null model are displayed in blue. Similarly, Fig. 4(c) shows the top features ranked according to statistical significance of feature coefficients. While caterpillar plots show the top-ranking features for a single value of *α*, eNetXplorer also generates heatmaps of feature frequencies (Fig. 4(b)) and feature coefficients (Fig. 4(d)) across all *α*–models.

**Figure 4:**
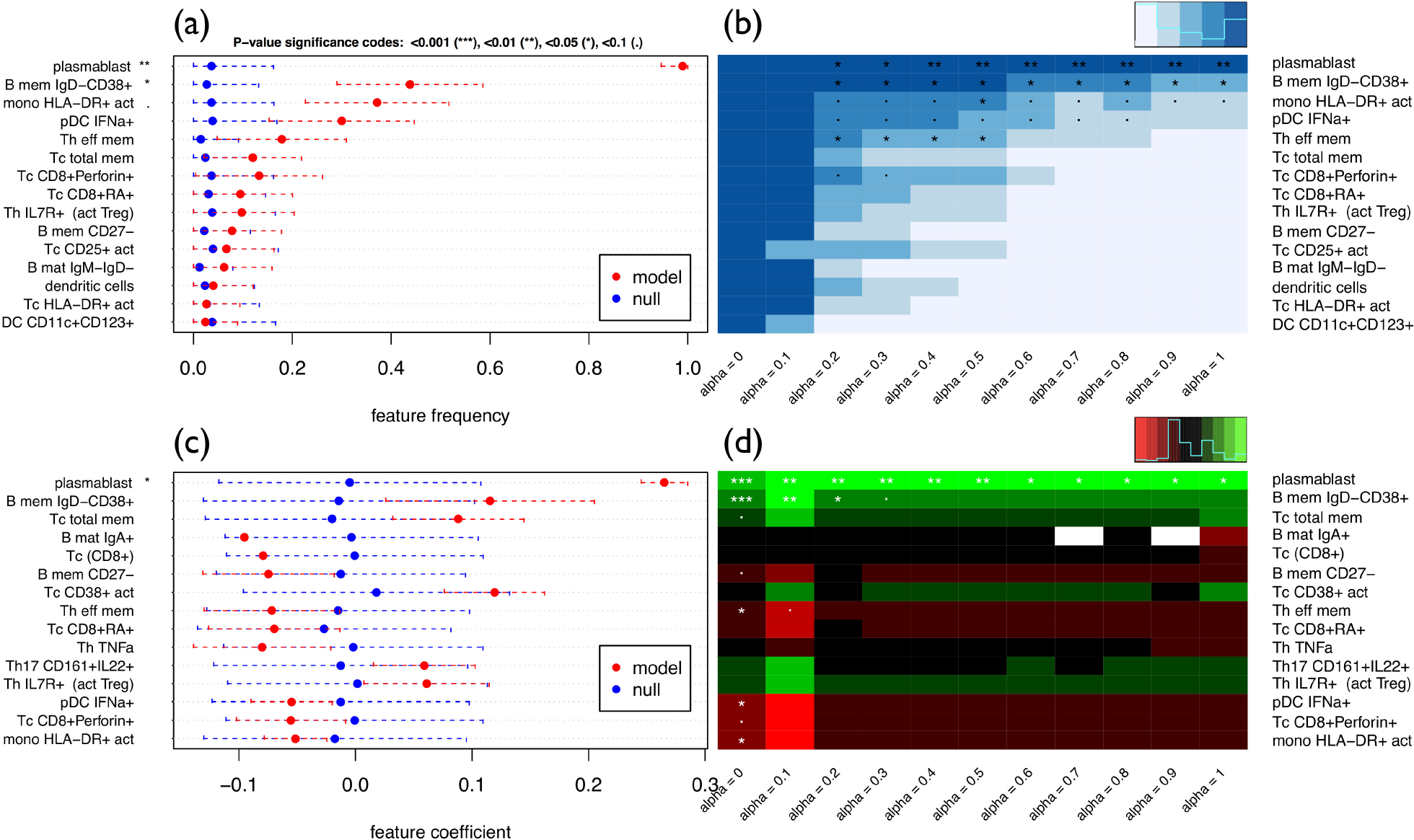
Linear regression model of postvaccination immune cell frequency correlates of H1N1 titer response. Top features selected from the lasso (*α* = 1) solutions. (a) Caterpillar plot of feature frequencies. Red symbols (bars) display the mean (standard deviation) of model feature frequencies across runs. Blue symbols (bars) display the mean (standard deviation) of null model feature frequencies across run/permutation null model combinations. (b) Heatmap of feature frequencies across *α*. (c) Caterpillar plot of feature coefficients. (d) Heatmap of feature coefficients across *α*. White denotes missing values, which occur for features whose frequencies are zero across all runs for a given *α*.

The same analysis strategy can be applied to any GLM in a similar fashion; two additional plot types are available to display results for binomial and multinomial classification models. Figs. 5(a,c) illustrate graphical representations of the contingency table for a multinomial classification analysis performed by eNetXplorer. Figs. 5(b,d) show boxplot representations of the OOB predictive accuracy for each class of samples, which can be thought of as the categorical counterparts of Fig. 3(c) for linear regression. Figs. 3-6 were generated by functions provided by eNetXplorer; these functions can be called with custom graphics parameters. The package also includes additional methods to provide summary and data export functionality to facilitate downstream analysis.

**Figure 5:**
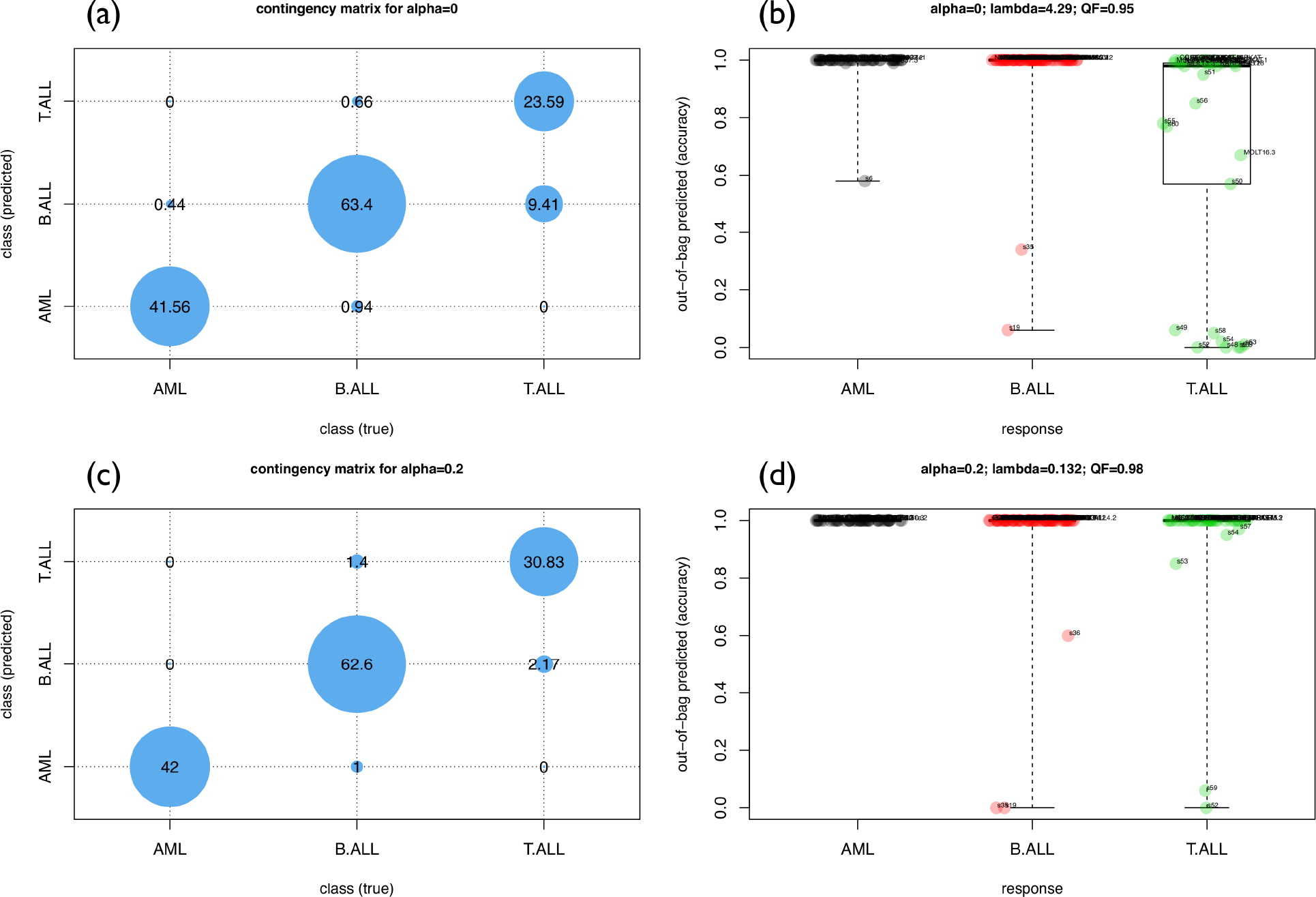
Multinomial classification model of microRNA expression in acute leukemias. Top (a,b): *α* = 0. Bottom (c,d): *α* = 0.2. Left (a,c): Contingency matrix of out-of-bag predictions vs response. Right (b,d): Boxplots of out-of-bag quality function (QF) per class across samples. The default QF for multinomial classification is the average accuracy, which is invariant under class label permutations.

## Results

### Linear regression case study: Predicting H1N1 influenza titers upon vaccination

Figs. 3-4 illustrate a typical eNetXplorer workflow to assess predictive models and parameters in the context of predicting antibody responses to H1N1 influenza vaccination using cell frequency data from Ref. ^7^ (included in the package). Here we focus on day 7 data (specifically, log fold-change from the baseline, i.e. log(day7)-log(day0)) to predict the antibody response on day 70.

The overall model performance across the entire elastic net family is summarized in Fig. 3(a), which shows that the statistical significance against the null model is *p* ∼ 0.1 for *α* ≃ 0.35 and increases monotonically towards the lasso (*p* ∼ 0.05). Based on the assessment for this specific dataset, we will focus our discussion on the lasso solution; however, it is useful to retain the ability to examine parameters for other values of *α*, which may pick up additional informative predictors and thus could provide further biological insights. It should be noted that the model-level statistical significance across *α* is dataset-specific; the accompanying vignette (Additional file 1) discusses varying effects of regularization on model performance, which arise in different scenarios of predictor/response covariance structure.

For *α* = 1, Fig. 3(b) shows the QF vs λ profile and the chosen λ* (dashed line) that was used to build the solution for this particular value of *α*. If, instead of displaying a well defined maximum, this profile happened to appear flat or monotonically increasing/decreasing, this could suggest that the range of λ is insufficiently large and needs to be extended via 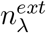, which is functionality implemented in eNetXplorer for this purpose. If the profile continues to appear flat or monotonic, that may suggest that the model is a poor fit to the data. Fig. 3(c) shows OOB predictions vs response for individual subjects. The positive correlation (*r* = 0.24) suggests that some cell populations at day 7 may indeed be informative of the antibody response after vaccination, although the substantial width of the 95% confidence interval (shown in light blue) suggests a weak statistical significance. This plot also highlights outliers such as subject ‘s244’, which appears with a large standard deviation and far from the region of correlation, which may be due to other covariates, such as demographic, clinical, or technical factors, not taken into account in the model. Note that in this illustrative analysis all subjects were used and the antibody response (as captured by the adjMFC metric ^7^) was modeled as a continuous variable in the linear regression. In the original publication, ^7^ the analysis focused on building predictive models and finding predictive parameters for high vs. low responders.

The caterpillar plot of Fig. 4(a) displays the top 15 cell populations ranked by model vs null significance according to feature frequency; the top features thus obtained are plasmablasts (*p* < 0.01) and IgD-CD38+ B-cell memory (*p* < 0.05), which were both reported as day 7 predictors in the original publication. ^7^ Fig. 4(b) shows these same top features (which were chosen based on the lasso solution) in the larger context of the entire elastic net family. Frequencies are trivially equal to 1 for ridge (*α* = 0), thus none appears as significant compared to the null; however, as *α* is increased, we observe the selection of several features that gradually decrease statistical significance towards the lasso. A complementary view is offered by the feature coefficient caterpillar plot (Fig. 4(c)) and corresponding heatmap (Fig. 4(d)), which show the direction (plus or minus sign) in which a given cell subpopulation affects the titer response. Taken together, feature frequency and feature coefficient maps point to potentially predictive cell populations, including those positively correlated with the titer response (plasmablasts, IgD-CD38+ memory B, CD25+ activated T cytotoxic) and others negatively associated with the titer response (HLA-DR+ activated monocytes, IFNa+ plasmacytoid dendritic cells, CD8+Perforin+ T cytotoxic, effector memory T helper).

## Multinomial classification case study: Uncovering microRNA

### signatures of acute leukemia subtypes

Figs. 5-6 present eNetXplorer results for multinomial models from a study of microRNA(miR)based signatures of acute myeloid leukemia (AML) in contrast to B-cell (B-ALL) and cell (T-ALL) lymphoblastic leukemias, ^8^ where features correspond to 370 miRs measured in multiple cell lines and primary leukemia samples. The full dataset is included in the package; details on data processing following Ref. ^8^ are provided in the accompanying vignette (Additional file 1).

Figs. 5(a,c) display the contingency matrix with the average number of instances predicted for each acute leukemia type; Figs. 5(b,d) show boxplot representations of OOB predicted samples in each class. The top panels correspond to ridge (*α* = 0), while the bottom panels correspond to slightly more regularized models (*α* = 0.2). The contingency matrix for ridge, Fig. 5(a), shows a good overall OOB classification performance, although with some misclassifications across the lymphoblastic classes; Fig. 5(b) displays predictions for individual samples. By increasing feature shrinkage, performance is quickly increased and the model is able to classify most samples correctly, as shown in Fig. 5(c-d). It is important to note that these results are based on a large number of cross validation iterations, where (for each run and for each fold within the run) a model was built using the in-bag, training data only, and the model was then applied to generate predictions on unseen, out-of-bag samples, followed by assessing the concordance between the predicted and known response (class labels). This process is free of data leakages since the training and testing sets are independent, thus mitigating the risk of having biased accuracy estimates due to overfitting.

For multinomial models, feature significance is separately assigned to each class. For AML, we observe that the top features selected by lasso are miR-27a, miR-223, and miR-145 (Fig. 6(a)), which agree with the most connected miRs in the cell-line based AML-centric dyad networks reported in Ref. ^8^ By considering less regularized models (i.e. for smaller values of *α*), miR-23a, miR-24, and other related miRs showed up as significantly associated with AML. miR-27a is co-localized in mammalian genomes with miR-23a and miR-24 and they form the so-called ‘miR-23a’ cluster, which was reported to be misregulated in multiple cancers; similarly, miR-145 was shown to be involved in proliferation and differentiation of hematopoietic cells and to be altered during leukemogenesis. ^9^ This example illustrates the ability of eNetXplorer to help explore AML signatures ranging from a minimal, informationally non-redundant set of markers (which can be useful prototypes of a diagnostic panel) to a larger, correlated set of signals (which can provide biological insight and guide the formulation of testable hypotheses). For B-ALL, we observe that miR-146a, miR-708, miR-629 and other significant miRs in the elastic net family (Fig. 6(c-d)) were also reported as hubs in B-ALL-centric network ensembles. ^8^ Most notably, quantitative reverse transcription polymerase chain reaction (qRT-PCR) analysis of relative miR-708 expression levels showed that it could be a good biomarker for B-ALL. ^8^ Similarly, the most significant, differentially expressed miRs for T-ALL previously reported are recapitulated, at various degrees of parsimony controlled by *α* (Fig. 6(e-f)). Beyond their potential role as diagnostic biomarkers, some of these differentially expressed microRNAs have been reported as clinically informative in the context of prognosis and treatment response in chronic and acute leukemia patients. ^10^

**Figure 6:**
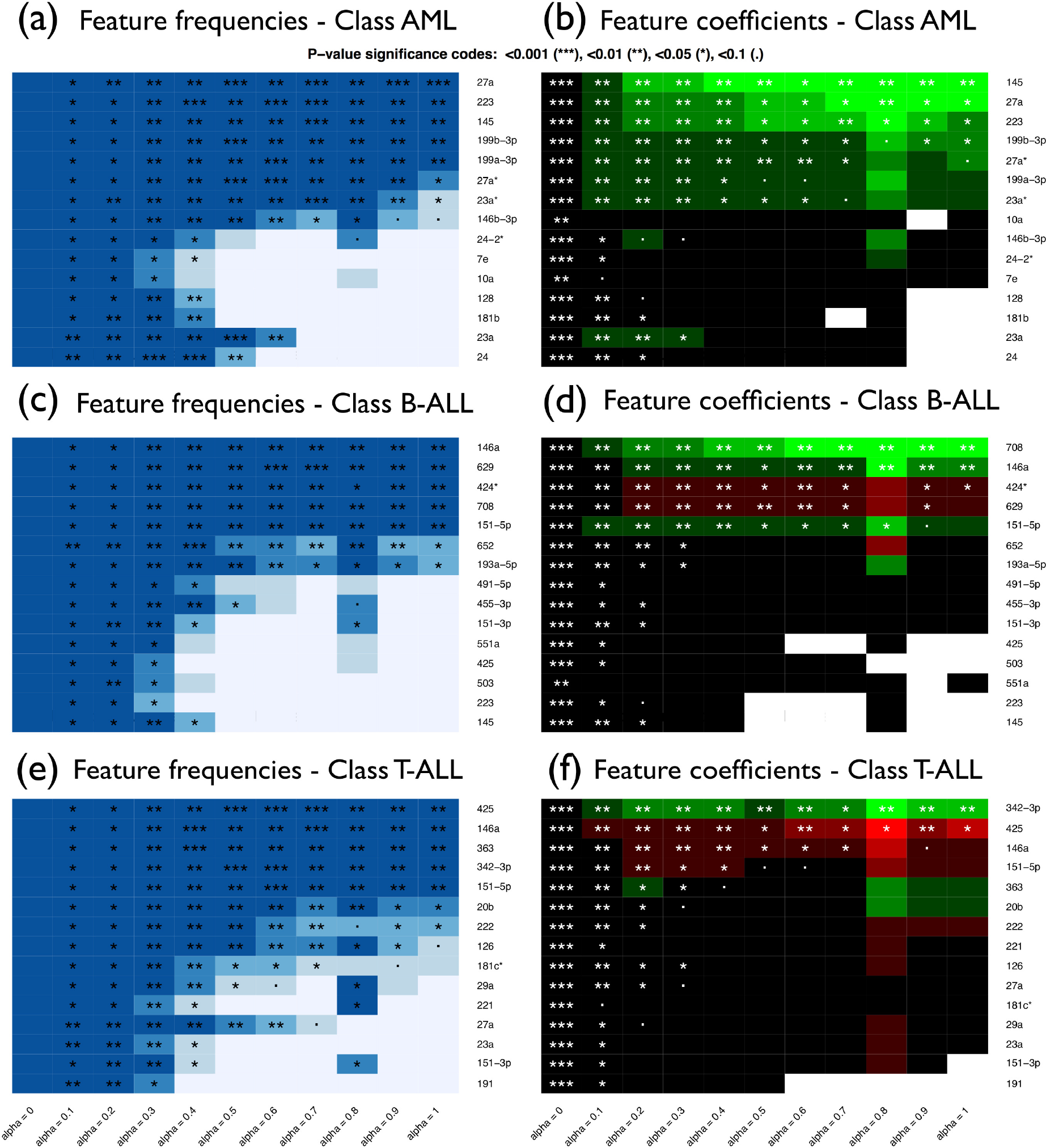
Multinomial classification model of microRNA expression in acute leukemias. Top features selected from the lasso (*α* = 1) solutions. Top (a,b): Acute myeloid leukemia (AML). Center (c,d): B-cell acute lymphoblastic leukemia (B-ALL). Bottom (e,f): T-cell acute lymphoblastic leukemia (T-ALL). Left (a,c,e): Heatmaps of feature frequencies. Right (b,d,f): Heatmaps of feature coefficients.

## Discussion

In a biomedical context, observations are associated with biological samples (derived from patients, model organisms, or cell lines), features are cellular and molecular measurements obtained from those samples (as well as demographic and clinical information associated with the subjects) and responses are categorical or numerical representations of phenotype, diagnosis, prognosis, or outcome (i.e. response to interventions). The number of available observations (*N*) in a study is severely constrained by limiting factors such as subject enrollment, ethical, financial and logistic considerations; on the contrary, the number of features per observation (*p*) enabled by state-of-the-art biotechnology assays and electronic health records is ever increasing. As represented by the conceptual schema of Fig. 1, regression models in these pervasive *p* ≫ *N* scenarios require regularization; the elastic net provides a framework to generate mixed-regularization model families. In this context, eNetXplorer plays a critical role by providing a quantitative assessment of model and feature performance, as well as of their statistical significance; it is to be viewed as the compass to navigate the regularization path.

In order to illustrate applications of eNetXplorer to real biomedical datasets, we presented two case studies. In the first one, we found cell populations that may explain the antibody response to H1N1 influenza vaccination. In the second study, we found micro-RNAs that may play key roles in leukemogenesis, and/or may be utilized as biomarker signatures. As discussed above, while some of the findings were validated by existing literature, others suggest novel associations that remain to be further explored. Naturally, regression models alone are unable to elucidate the molecular mechanisms at play; their role is that of showing (potentially novel) associations in large and complex datasets, thus aiding field experts in the process of formulating hypotheses and suggesting further experiments to confirm or rule out those hypotheses.

Lastly, let us emphasize that eNetXplorer, although primarily conceived in the context of biomedical research, it is generally applicable to other research areas as well. The accompanying vignette (Additional file 1) illustrates eNetXplorer workflows of general applicability.

## Conclusions

Uncovering correlates and predictors in a multi-parameter setting is an ubiquitous problem in systems biology. In this context, regularized generalized linear modeling is a popular approach due to its flexibility, but it is often desirable to retain the ability to explore different levels of regularization and examine elastic net families that span the full range from ridge to lasso.

Our package is built on top of glmnet to provide novel functionalities that neither glmnet itself, nor (to the best of our knowledge) other currently available software packages provide. Importantly, one of the most valuable new functionalities our software enables is to empower biological applications in real-world settings to address one of the most frequently asked questions: which biological variables, often out of a large number in the current age of large-scale ‘omicsâĂŹ datasets, provide predictive information about an outcome variable (e.g. diagnosis, vaccination efficacy, drug/treatment response, etc.)? Specifically, both the null model evaluation functions (based on response label permutations) that quantitatively assess which parameters are important and statistically significant for prediction, as well as a set of functions for visualization of these results across parameter space provided by our software, are novel and provide a systematic framework to rigorously assess parameter significance.

Thus, eNetXplorer aims to make regularization approaches to generalized linear modeling more readily available to a larger user base of diverse scientific background in order to transform large-scale data sets into biological hypotheses and insight.

## Availability and requirements

Project name: eNetXplorer

Project home page: https://CRAN.R-project.org/package=eNetXplorer

Operating system(s): Platform independent Programming language: R (≥ 2.10)

Other requirements: R packages glmnet, stats, Matrix, RColorBrewer, calibrate, progress, graphics, methods, grDevices, gplots

License: GPL-3

Any restrictions to use by non-academics: none

## List of abbreviations

GLM: Generalized linear model
QF: Quality function
OOB: Out-of-bag
miR: micro-RNA
AML: Acute Myeloid Leukemia
B-ALL: B-cell Acute Lymphoblastic Leukemia
T-ALL: T-cell Acute Lymphoblastic Leukemia

## Declarations

### Ethics approval and consent to participate

Human data described in this paper belong to the public domain. The H1N1 influenza study was originally published in Ref.; ^7^ this investigation was approved by the Institutional Review Board of the National Institutes of Health (Protocol 09-H-0239; ClinicalTrials.gov Identifier: NCT01191853) and conducted according to the principles expressed in the Declaration of Helsinki. The acute leukemia study was originally published in Ref.; ^8^ this investigation was approved by the Institutional Review Board of the University of Maryland Baltimore and conducted according to the principles expressed in the Declaration of Helsinki.

### Consent for publication

Not applicable.

### Availability of data and materials

The eNetXplorer R package is available under GPL-3 license at the CRAN repository, https://CRAN.R-project.org/package=eNetXplorer. The datasets discussed here are distributed with the package.

### Authors’ contributions

JC designed the project, implemented the code in R, wrote the paper. JST designed the project, wrote the paper. All authors discussed the manuscript, read and approved the final version of the manuscript.

## Acknowledgements

The authors thank Angelique Biancotto, Jinguo Chen, Foo Cheung, Yuri Kotliarov, and Pamela Schwartzberg for discussions and feedback during the development of this package.

## Additional files

Additional file 1: eNetXplorer vignette. Detailed description of eNetXplorer’s workflow applied to synthetic datasets with different feature/response covariance structures. Descriptions of real datasets distributed with the package and analyzed in the Results Section of this paper are also provided, as well as a Summary with the key aspects of eNetXplorer. (PDF 325 kb)

