## Additional file 1 for "eNetXplorer: an R package for the quantitative exploration of elastic net families for generalized linear models"

### eNetXplorer Vignette

#### Table of contents

1. About
2. Installation
3. eNetXplorer's Workflow
4. Datasets
    1. `H5N1_Flow`
    2. `Leukemia_miR`
5. Summary

#### 1. About

The R package `eNetXplorer` is available under GPL-3 license at the CRAN repository. The package source is located at https://CRAN.R-project.org/package=eNetXplorer.

Authors: Julián Candia and John S. Tsang

Maintainer: Julián Candia <>

#### 2. Installation

To install to your default directory, type

```
install.packages("eNetXplorer")
```

For more installation options, type `help(install.packages)`.

#### 3. eNetXplorer's Workflow

In order to describe `eNetXplorer`'s workflow, this Section presents the analysis pipeline applied to synthetic datasets; to further illustrate eNetXplorer's features, real datasets are distributed with the package as described in Sections below.

First, we create a function to generate data for a Gaussian (linear regression) model, which consists of:

- An input numerical matrix with `n_inst` instances or observations (as rows) and `n_pred` predictors or features (as columns).
- An input numerical vector of length `n_inst` with the observed response.

For normally distributed random variables, the following data-generating function will create a set of features correlated with the response according to any arbitrary, pre-defined population covariance matrix `covmat`:

```
data_gen <- function(n_inst, covmat, seed=123) {
    library (expm);
    set.seed(seed)
    data <- matrix(rnorm(n_inst*ncol(covmat)),ncol=ncol(covmat))%*%sqrtm(covmat)
    predictor=data[,-1]
    rownames(predictor) = paste0("Inst.",1:n_inst)
```

```
    colnames(predictor) = paste0("Feat.",1:(ncol(covmat)-1))
    list(response=data[,1],predictor=predictor)
}
```

Here, we assume that the first row/column in `covmat` is the response, while the remaining variables correspond to the features. In order to generate covariant matrices with interesting properties, the following function will produce a block of features correlated with each other (where `r_block` is the intra-block Pearson's correlation) as well as correlated with the response (with correlation `r_resp`):

```
covmat_gen <- function(n_pred, block_size, r_resp, r_block) {
    covmat = matrix(rep(1.e-3,(n_pred+1)**2),ncol=(n_pred+1))
    for (i_pred in 1:block_size) {
        for (j_pred in (i_pred+1):(block_size+1)) {
            if (i_pred==1) {
                covmat[i_pred,j_pred] = r_resp
            } else {
                covmat[i_pred,j_pred] = r_block
            }
            covmat[j_pred,i_pred] = covmat[i_pred,j_pred]
        }
    }
    for (i_pred in 1:n_pred) {
        covmat[i_pred,i_pred] = 1
    }
    covmat
}
```

Let us use these two functions to generate predictor and response inputs:

```
data = data_gen(n_inst=50, covmat_gen(n_pred=60, block_size=5, r_resp=0.5, r_block=0.35))
```

We examine the predictor correlation structure; as expected from our `covmat_gen` call, the block of features 1-5 is significantly intra-correlated, while the remaining features appear only weakly (and, in some cases, even negatively) correlated.

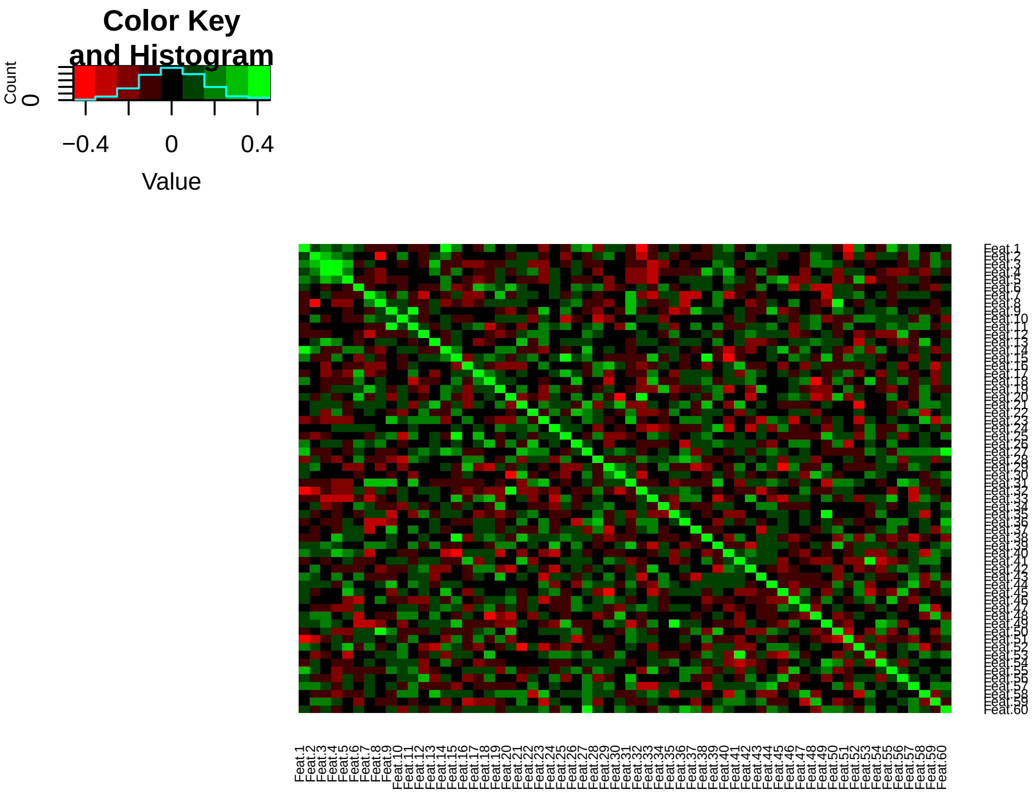


We also examine the correlation between predictors and the response; by design, the block of features 1-5 carries the largest positive correlation with the response.

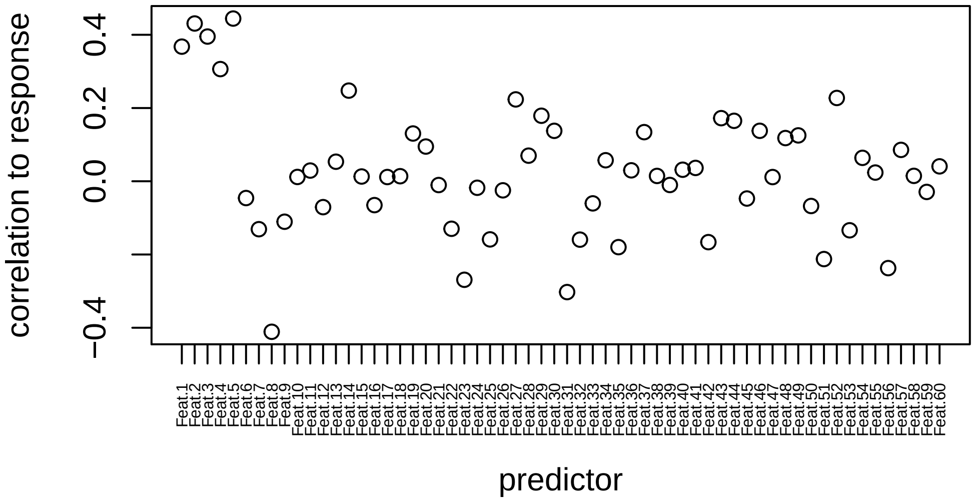


Since the number of features is larger than the number of observations, we need to implement a regularized regression model. But which model? We know that ridge will fit a model where all predictors (including all the non-informative ones) will have non-zero contributions. Lasso, on the other end, will exclude most features and provide a minimal model representation. The elastic net allows us to scan the regularization path from ridge to lasso via the mixing parameter `alpha`. However, the following open questions remain:

- Which `alpha` represents the top-performing regularized model?
- What is the model-level statistical significance across `alpha`?
- What is the feature-level statistical significance of a given `alpha`-model?
- How does feature-level statistical significance change across `alpha`?

To address these questions, `eNetXplorer` generates an ensemble of null models (based on random permutations of the response) on a family of regularized models from ridge to lasso. First, we load the `eNetXplorer` package:

```
library(eNetXplorer)
```

Next, we run `eNetXplorer` on the datasets we just generated. The call to `eNetXplorer` with default parameters is:

```
fit_def = eNetXplorer(x = data$predictor, y = data$response, family = "gaussian")
```

Results can be made more precise by increasing the number of cross-validation runs (`n_run`) and the number of null-model response permutations per run (`n_perm_null`), as well as by choosing a smaller step in the path of `alpha` models:

```
fit = eNetXplorer(x = data$predictor, y = data$response, family = "gaussian",
    alpha = seq(0, 1, by = 0.1), n_run = 1000, n_perm_null = 250, seed = 123)
```

Function `summary` generates a brief report on the results; for each `alpha`, it displays the optimal `lambda` (obtained by maximizing a quality function over out-of-bag instances), the corresponding maximum value of the quality function, and the model significance (p-value based on comparison to permutation null models).

```
summary(fit)
```

```
## Call:
## eNetXplorer(x = data$predictor, y = data$response, family = "gaussian",
##     alpha = seq(0, 1, by = 0.1), seed = 123, n_run = 1000, n_perm_null = 250)
##
##     alpha lambda.max QF.est model.vs.null.pval
##  0.000000   3.724056 0.2222            0.06524 .
##  0.100000   0.697820 0.3202            0.04222 *
##  0.200000   0.382928 0.3254            0.04641 *
##  0.300000   0.293516 0.3096            0.05316 .
##  0.400000   0.200581 0.3086            0.06305 .
##  0.500000   0.168105 0.3038            0.06389 .
##  0.600000   0.140088 0.2936            0.07106 .
##  0.700000   0.125793 0.2936            0.07027 .
##  0.800000   0.115310 0.2923            0.07448 .
##  0.900000   0.097839 0.2886            0.07634 .
##  1.000000   0.092248 0.2776            0.08398 .
## ---
## Signif. codes:  0 '***' 0.001 '**' 0.01 '*' 0.05 '.' 0.1 ' ' 1
```

A graphical display of model performance across `alpha` is provided by

```
plot(fit, plot.type="summary")
```

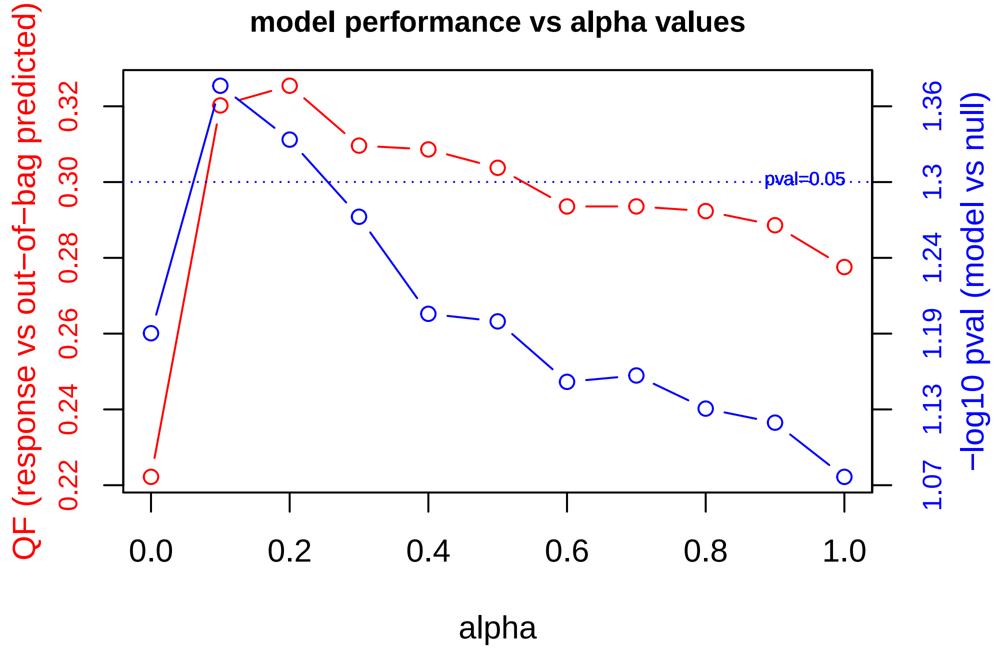


We observe that, at `p-value`<0.05, the most significant models are `alpha`=0.1 and 0.2; in terms of performance, the quality function (which, by default, is Pearson's correlation) evaluated between out-of-bag predictions and the response is maximized by the `alpha`=0.2 model. Let us examine this top-performing model in more detail.

Our next question is to determine the top features that play a role in the top-performing model. Following a similar strategy to that of `alpha`-model selection, statistical significance at the feature level is determined by comparison to permutation null models; see (Candia and Tsang 2019) for technical details.

We generate a caterpillar plot of the top features based on their coefficients:

```
plot(fit, alpha.index = which.max(fit$model_QF_est), plot.type = "featureCaterpillar",
    stat = c("coef"))
```

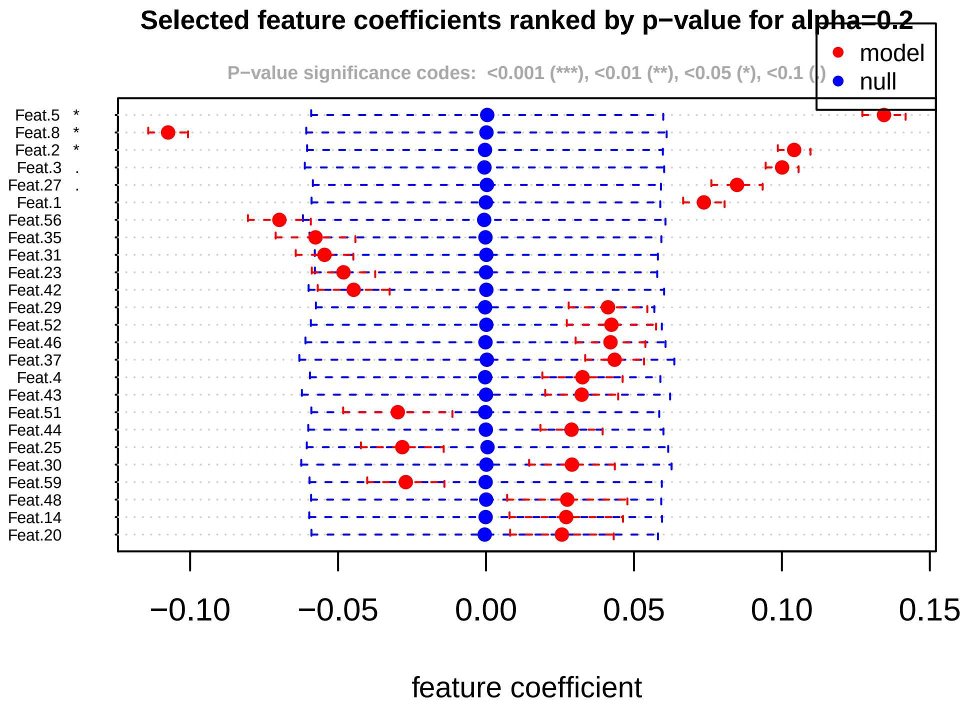


We observe that features 5, 8, and 2 are selected by the regularized model with `alpha`=0.2 at `p-value`<0.05; features 3 and 27 are significant at `p-value`<0.1. Next, we aim to explore those same top features across the entire elastic net family: Which of them would still be selected under more stringent regularization criteria?

```
plot(fit, alpha.index = which.max(fit$model_QF_est), plot.type = "featureHeatmap",
    stat = c("coef"), notecex = 1.5)
```

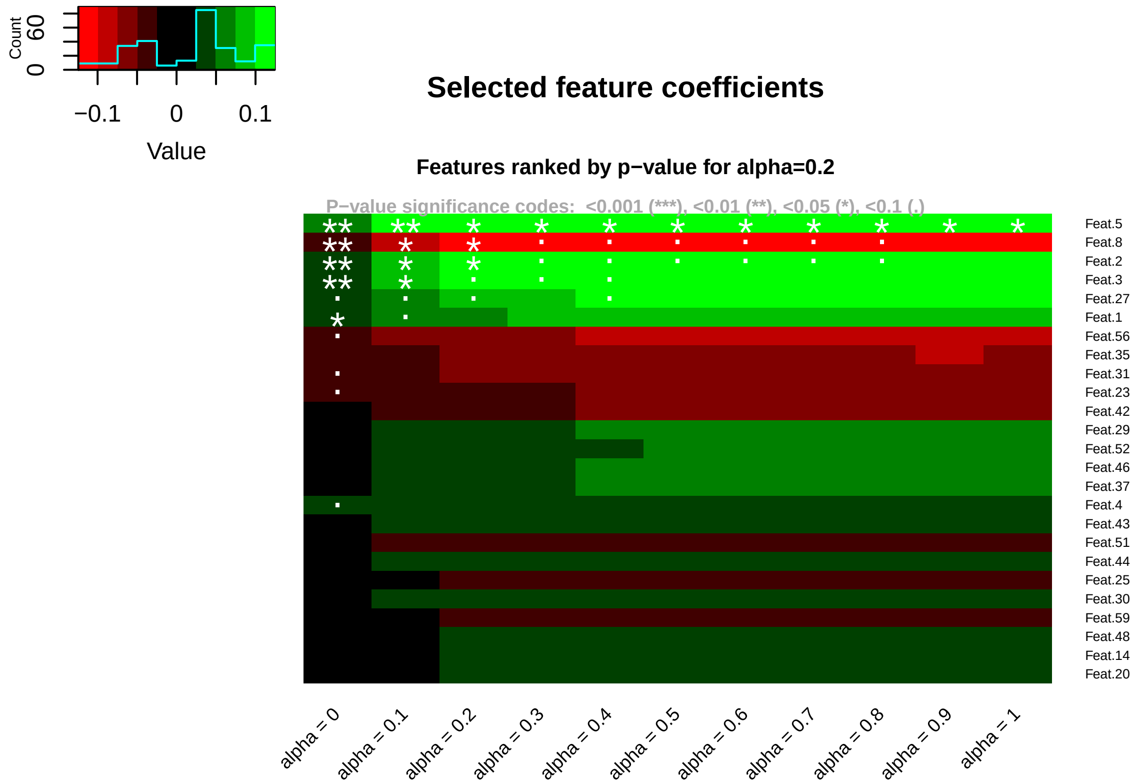

Here, we observe that more regularized models (`alpha`=0.5-0.8) favor features 5, 8 and 2 (at significance `p-value`<0.1); towards lasso (`alpha`=0.9,1), feature 5 is the only one selected at `p-value`<0.05-0.1. As expected, these models are more stringent on feature selection than less regularized (i.e. smaller `alpha`) models. It is also interesting to observe that lasso-like models remove feature redundancies: from the block of correlated features 1-5, only feature 5 is picked up as representative. This example illustrates some important characteristics of mixed-regularization model families:

- less regularized (smaller `alpha`) models promote redundancy; they benefit from borrowed information across significantly correlated predictors; they provide larger signatures, which are potentially more robust and resilient under measurement noise; they offer more opportunities for systems-level interpretation (e.g. downstream pathway analysis in the context of genomics).
- more regularized (larger `alpha`) models promote sparsity; they tend to pick just one predictor out of a set of correlated ones; they may facilitate interpretation with high-dimensional datasets and/or in the absence of systems-level annotations; they provide smaller signatures, which may be more useful in certain contexts (e.g. biomarker panels).

In order to gather more details regarding a particular solution, we plot the quality function across the range of values for the regularization parameter `lambda`:

```
plot(fit, alpha.index = which.max(fit$model_QF_est), plot.type = "lambdaVsQF")
```

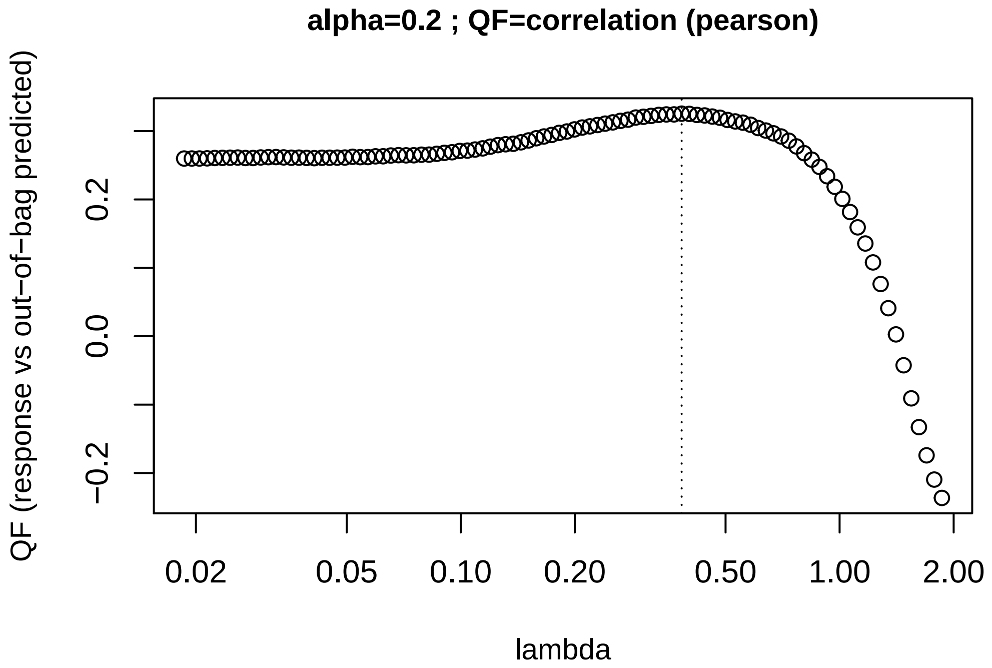


If so desired, `eNetXplorer` allows end-users to extend the number of `lambda` values (via `nlambda`) and/or extend their range while keeping the `lambda` density uniform in log scale (via `nlambda.ext`).

There may exist outlier instances that may require further examination; we generate a scatterplot of response vs out-of-bag predictions across all instances:

```
plot(fit, alpha.index = which.max(fit$model_QF_est), plot.type = "measuredVsOOB")
```

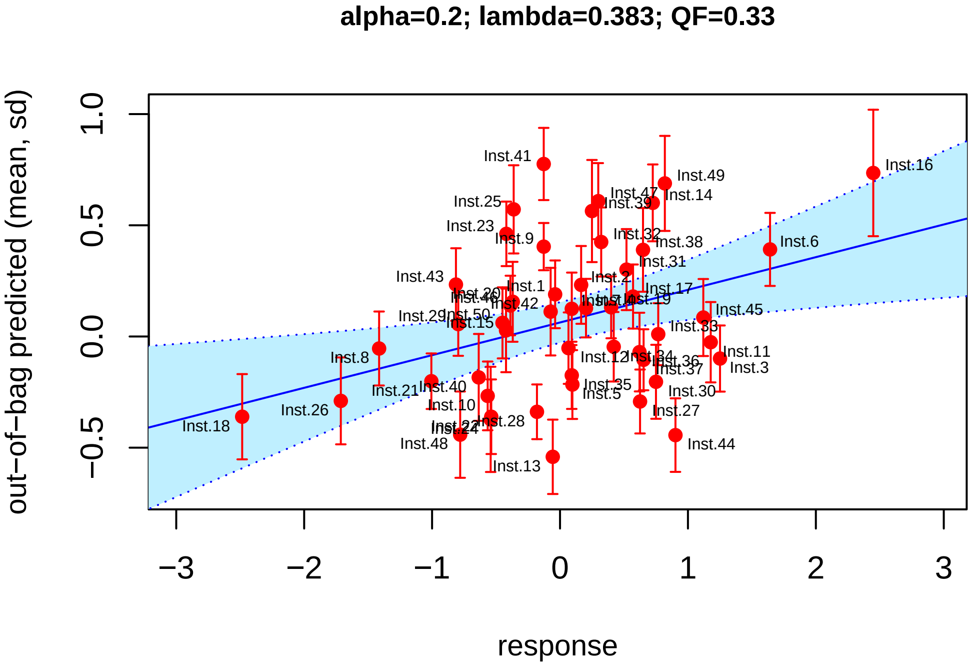


Naturally, model performance across `alpha` is strongly dependent on the structure of the input datasets. In the case example above, we purposefully generated a block of informative correlated predictors to highlight the characteristics of less regularized (smaller `alpha`) models and their ability to leverage borrowed information. Let us now generate input datasets with just one prevalent informative predictor:

```
data = data_gen(n_inst=50, covmat_gen(n_pred=60, block_size=1, r_resp=0.7, r_block=0.35))
```

As before, we examine the predictor correlation structure:

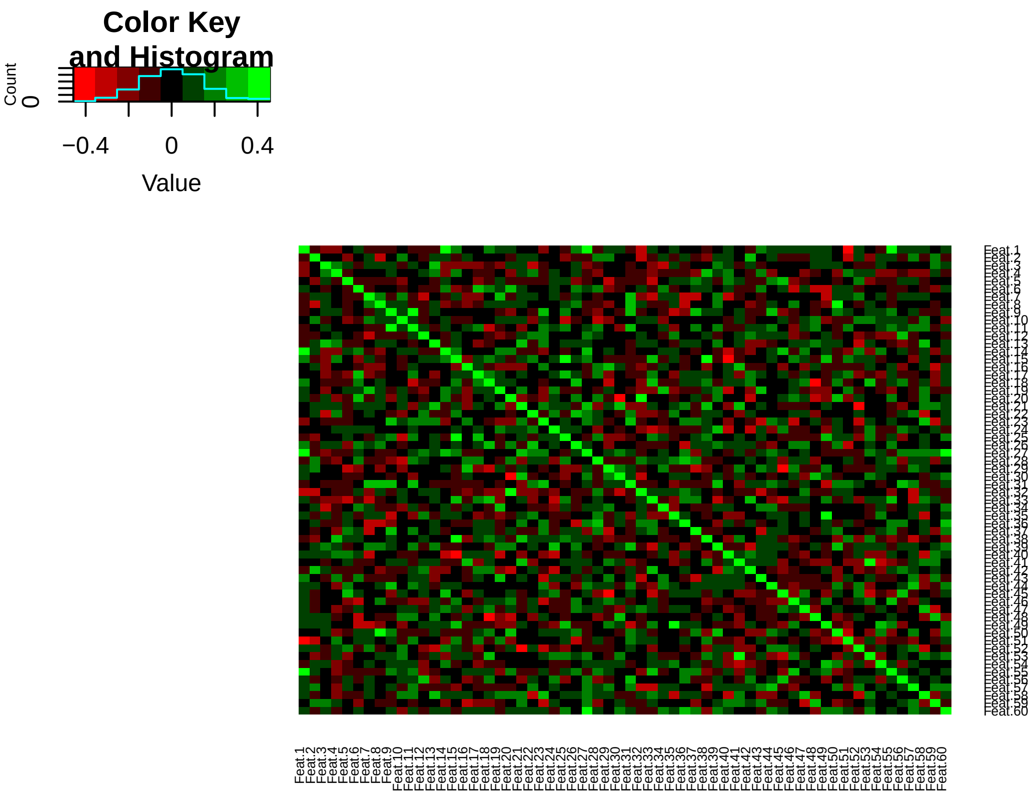

Moreover, as before, we also examine the correlation between predictors and the response; by design, feature 1 is the most informative predictor:

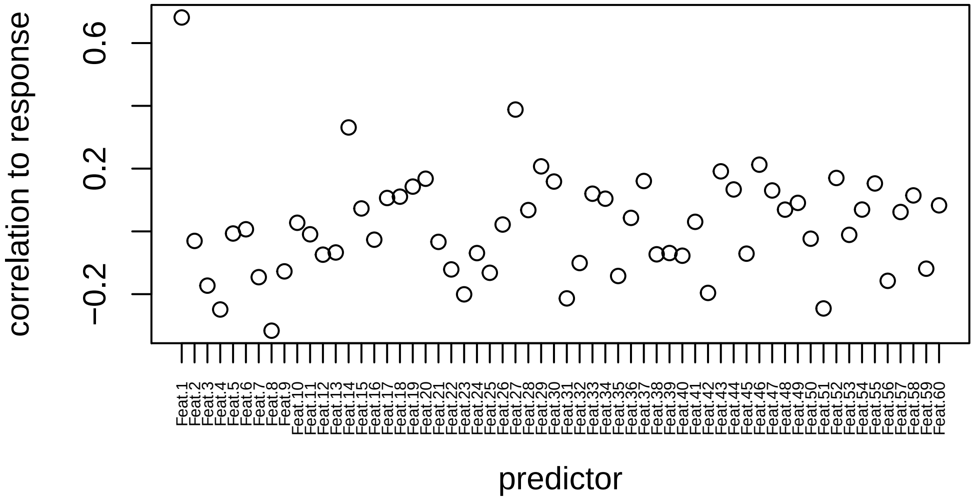


We run `eNetXplorer` on the new datasets:

```
fit = eNetXplorer(x = data$predictor, y = data$response, family = "gaussian",
    alpha = seq(0, 1, by = 0.1), n_run = 1000, n_perm_null = 250, seed = 123)
```

The summary table is:

```
summary(fit)
```

```
## Call:
## eNetXplorer(x = data$predictor, y = data$response, family = "gaussian",
##     alpha = seq(0, 1, by = 0.1), seed = 123, n_run = 1000, n_perm_null = 250)
##
##    alpha lambda.max QF.est model.vs.null.pval
##  0.00000    6.15154 0.1833           0.082884 .
##  0.10000    1.92280 0.4391           0.004960 **
##  0.20000    1.21315 0.5179           0.000824 ***
##  0.30000    0.88762 0.5475           0.000384 ***
##  0.40000    0.66571 0.5691           0.000180 ***
##  0.50000    0.53257 0.5824           0.000156 ***
##  0.60000    0.42364 0.5916            8.0e-05 ***
##  0.70000    0.38041 0.5950           0.000116 ***
##  0.80000    0.33286 0.6026            8.8e-05 ***
##  0.90000    0.29587 0.6042           0.000108 ***
##  1.00000    0.27896 0.6094            8.0e-05 ***
## ---
## Signif. codes:  0 '***' 0.001 '**' 0.01 '*' 0.05 '.' 0.1 ' ' 1
```

We generate the plot of model performance across `alpha`:

```
plot(fit, plot.type="summary")
```

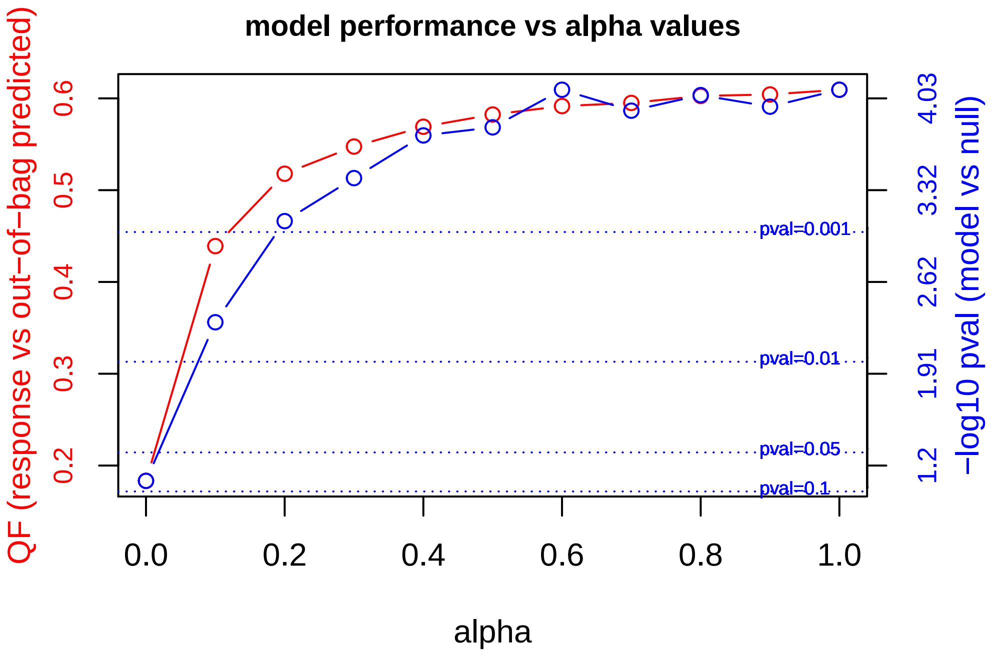


Here, we observe that the quality function appears to increase monotonically with `alpha`; the maximum corresponds to the lasso solution, `alpha`=1.

We generate a caterpillar plot of the top features based on their coefficients:

```
plot(fit, alpha.index = which.max(fit$model_QF_est), plot.type = "featureCaterpillar",
    stat = c("coef"))
```

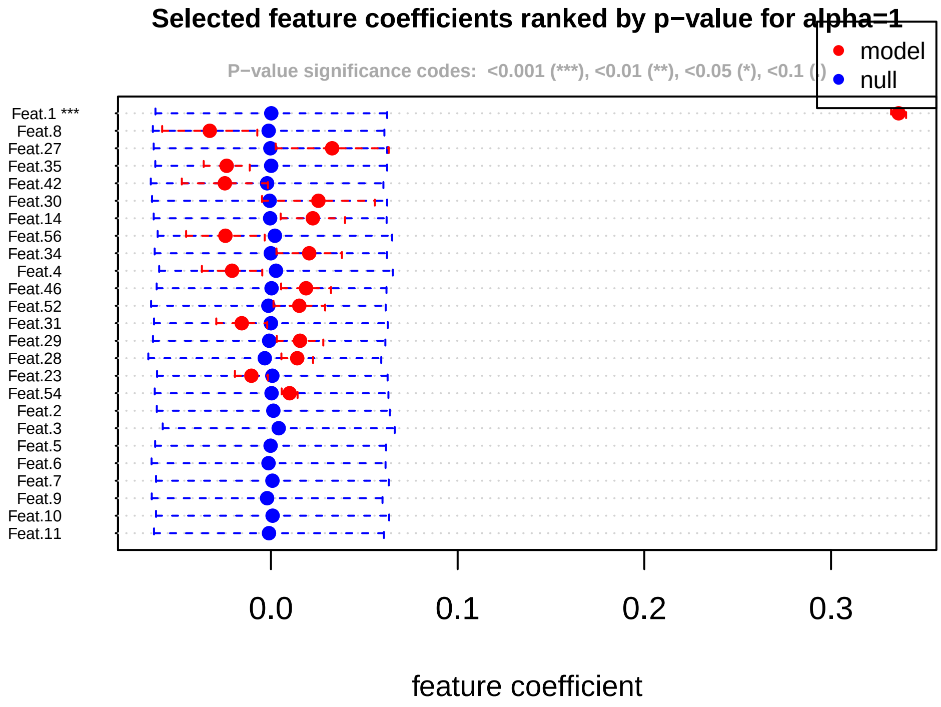


We observe that only feature 1 is selected by lasso. Are there other features that would be selected under less stringent regularization criteria?

```r
plot(fit, alpha.index = which.max(fit$model_QF_est), plot.type = "featureHeatmap",
    stat = c("coef"), notecex = 1.5)
```

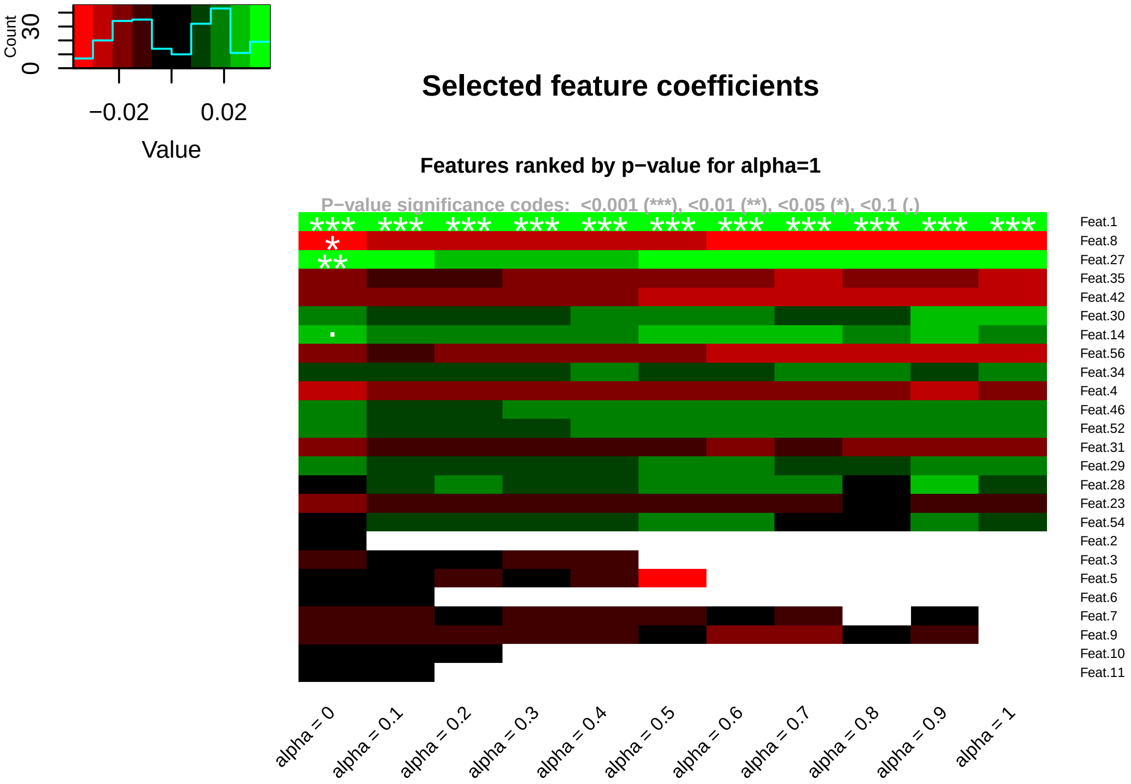


We find that, except for features 8, 27 and 14 for ridge regularization (`alpha`=0), feature 1 appears dominant all across the regularization range. This, indeed, reflects the covariance structure we chose to generate the data.

### 4. Datasets

The `eNetXplorer` package provides two datasets of biological interest:

- `H1N1_Flow`, comprised of longitudinal cell population frequencies and titer responses upon H1N1 vaccination; and
- `Leukemia_miR`, which contains microRNA (miR) expression data from cell lines and primary (patient) samples classified by different acute leukemia phenotypes, as well as normal control samples sorted by cell type.

#### 4.1 `H1N1_Flow`

The `H1N1_Flow` dataset comprises data from a cohort of healthy subjects vaccinated against the influenza virus H1N1 (Tsang et al. 2014). Using five different 15-color flow cytometry stains for T-cell, B-cell, dendritic cell, and monocyte deep-phenotyping, 113 cell population frequencies were measured longitudinally pre- (days -7, 0) and post-vaccination (days 1, 7, 70) on a cohort of 49 healthy human subjects (F=31, M=18, median age=24). Cell populations were manually gated and expressed as percent of parent. Samples and cell populations were filtered independently for each timepoint; samples were excluded if the median of the fraction of viable cells across all five tubes was <0.7, while cell populations were excluded if >80% of samples had <20 cells. Data were log10-transformed and pooled across all timepoints, then adjusted for age, gender, and ethnicity effects. For each timepoint, a numerical matrix of predictors is provided with subjects as rows and cell populations as columns. The response is the adjusted maximum fold change (adjMFC) of serum titers at day 70 relative to baseline, as defined in (Tsang et al. 2014). Two versions of the serum titer response are provided in the package; one as a numerical vector and the other one as a categorical vector discretized into low ("0"), intermediate ("1") and high ("2") response classes. A metadata file with cell population annotations is also provided.

To load the dataset:

```
data(H1N1_Flow)
```

##### 4.2 Leukemia_miR

The `Leukemia_miR` dataset comprises data of human microRNA (miR) expression of 847 miRs from 80 acute myeloid (AML) and acute lymphoblastic (ALL) leukemia cell lines, 60 primary (patient) samples, and 50 normal control samples sorted by cell type (CD34+ HSPC, Granulocytes, Monocytes, T-cells and B-cells) (Tan et al. 2014; Candia et al. 2015). Acute lymphoblastic leukemia samples are further classified by B-cell (B-ALL) and T-cell (T-ALL) subphenotypes. Two dataset versions are provided: the full dataset `Leuk_miR_full` (190 samples x 847 miRs) and the filtered dataset `Leuk_miR_filt` (140 samples x 370 miRs). A numerical matrix of predictors is provided with samples as rows and miRs as columns. Two categorical response vectors are provided for binomial (AML, ALL) and multinomial (AML, B-ALL, T-ALL) classification.

To load the dataset:

```
data(Leukemia_miR)
```

To filter the full dataset:

```
expr_full = Leuk_miR_full$expression_matrix
miR_filter = rep(F,nrow(Leuk_miR_full$miR_metadata))
miR_filter[apply(expr_full,2,mean)>1.2] = T
sample_filter = rep(T,nrow(Leuk_miR_full$sample_metadata))
sample_filter[Leuk_miR_full$sample_metadata$sample_class=="Normal"] = F
expr_filtered = expr_full[sample_filter,miR_filter]
miR_filtered = Leuk_miR_full$miR_metadata[miR_filter,]
sample_filtered = Leuk_miR_full$sample_metadata[sample_filter,]
```

#### 5. Summary

- `eNetXplorer` addresses key questions regarding model regularization and feature selection based on permutation-based statistical significance tests. Results are provided in the form of standard plots, summary statistics and output tables.
- As illustrated by synthetic datasets, which were generated from covariance matrices involving normally distributed features and response distributions, `eNetXplorer` workflows are generally applicable to model any datasets consisting of `n_inst` observations across `n_pred` predictors or features that aim to explain a set of `n_inst` response values.
- Regularization models are particularly useful in scenarios involving a large number of features (even larger than the number of observations) and/or sets of significantly correlated features. These scenarios are typical of datasets generated by current technologies in molecular and cellular biology, but applications to other data rich environments are certainly possible.
